# JG2: an updated version of the Japanese population-specific reference genome

**DOI:** 10.1101/2024.11.01.621223

**Authors:** Sirawit Sriwichaiin, Satoshi Makino, Takamitsu Funayama, Akihito Otsuki, Junko Kawashima, Yasunobu Okamura, Shu Tadaka, the Tohoku Medical Megabank Project Study Group, Fumiki Katsuoka, Kazuki Kumada, Shuichi Tsutsumi, Kengo Kinoshita, Masayuki Yamamoto, Gen Tamiya, Jun Takayama

**Affiliations:** Department of AI and Innovative Medicine, Tohoku University School of Medicine, 2-1, Seiryo-machi, Aoba-ku, Sendai, Miyagi 980-8573, Japan; Tohoku Medical Megabank Organization, Tohoku University, 2-1, Seiryo-machi, Aoba-ku, Sendai, Miyagi 980-8573, Japan; Advanced Research Center for Innovations in Next-Generation Medicine, Tohoku University, 2-1 Seiryo-machi, Aoba-ku, Sendai, Miyagi 980-8573, Japan; Genome Science Division, Research Center for Advanced Science and Technology, The University of Tokyo, Tokyo, Japan; Department of Applied Information Sciences, Graduate School of Information Sciences, Tohoku University, Sendai, Miyagi 980-8579, Japan; Department of In Silico Analyses, Institute of Development, Aging and Cancer (IDAC), Tohoku University, Sendai, Miyagi 980-8575, Japan; Department of Biochemistry and Molecular Biology, Tohoku Medical Megabank Organization, Tohoku University, 2-1 Seiryo-machi, Aoba-ku, Sendai, Miyagi 980-8573, Japan; Statistical Genetics Team, RIKEN Center for Advanced Intelligence Project, Nihonbashi 1-chome Mitsui Building 15F, 1-4-1 Nihonbashi, Chuo-ku, Tokyo 103-0027, Japan

## Abstract

This study presents the construction of JG2, an updated population-specific reference genome for the Japanese population. Utilizing data from three individuals previously employed in the construction of JG1, several methodologies were employed to enhance genomic coverage and assembly quality. Hi-C sequencing technology facilitated phase-aware assembly, generating two haploid assemblies per individual and enabling improved representation of genetic variation. A meta-assembly strategy and a majority decision approach further refined assembly quality by combining the best sequences from multiple assemblies and minimizing the inclusion of rare variants. The resulting JG2 genome comprises chromosome-level sequences, mitochondrial chromosomes, and unplaced scaffolds, offering more comprehensive coverage of the Japanese genome. Comparative analyses with other reference genomes demonstrated the accuracy and representativeness of JG2, highlighting its utility for genetic research involving the Japanese population. Overall, by adopting the phased assembly technique, JG2 represents a significant advancement over the collapsed assembly-based JG1, providing researchers with a more precise and comprehensive resource for understanding the genetic landscape of the Japanese population. The sequences and annotations are available on the jMorp website (https://jmorp.megabank.tohoku.ac.jp/).

## Introduction

Despite the development of human reference genomes (1, 2), population-specific reference genomes are crucial for accurately capturing the genetic diversity and unique variations within distinct human populations. Several population-specific reference genomes were constructed and demonstrated novel genetic diversity in a population-specific manner (3–8). To address this need, we previously developed JG1, a population-specific reference genome for the Japanese population (8). JG1 incorporates the major alleles of Japanese individuals as reference alleles for variants, including single nucleotide variants (SNVs), short insertion and deletions (indels), and structural variants (SVs). This was achieved by integrating *de novo* assemblies from three Japanese individuals, using meta-assembly strategies and majority decision among multiple genome assemblies to reduce the impact of rare or private variants. Additionally, by anchoring scaffolds with marker information from genetic and radiation hybrid maps, JG1 was constructed independently of other human reference genomes, marking a significant milestone in creating a *de novo* Japanese reference genome. JG1 is also used for NGS analyses to identify the causal variants of rare diseases in several studies (9–12).

Despite its successes, JG1 has limitations, such as incomplete sequences, gaps, unlocalized fragments, limited original annotations, and an incomplete representation of major variations within the Japanese population. Addressing these limitations could significantly enhance the quality of the human reference genome, especially benefiting genome research involving Japanese or Asian populations. Thus, we developed JG2, an updated version of JG1, to overcome these challenges.

The construction of JG2, like JG1, was independent of the other human reference genomes. We utilized phased assembly, employing Falcon, Falcon Unzip (13), and Falcon Phase algorithms with Hi-C reads (14) on Pacific Biosciences (PacBio) RS II continuous long reads (CLR) from the three individuals. Hi-C reads enabled the integration of long-range chromatin interaction data, essential for resolving complex genomic regions and producing a more precise and continuous genome assembly (15). This method is particularly beneficial for phased assembly, as it aids in differentiating between maternal and paternal haplotypes. Therefore, this approach resulted in two sets of haploid assembly per individual. The scaffolding process of fully phased contigs was also conducted by incorporating information from Hi-C reads, yielding six PacBio/Hi-C phased scaffolds. These six haploid scaffolds were then subjected to a meta-assembly process, leveraging the concept that two haploid genomes represent a random sample from a population, enhancing the representativeness of genetic variations. Building upon the techniques employed in the construction of JG1, a meta-assembly was utilized to reconcile multiple assemblies. Scaffolds were anchored using marker information from genetic and radiation hybrid maps, ensuring that JG2’s reconstruction remained independent of other reference genomes. A majority decision of alleles was also conducted based on six haploid scaffolds. This process reduced the rare variants in JG2, making it more representative of the general Japanese population. The genome sequences and annotations of JG2 are available on the jMorp website (https://jmorp.megabank.tohoku.ac.jp/).

## Materials and Methods

### Ethics declaration

This study received approval from the Research Ethical Committee of the Tohoku Medical Megabank Organization, Tohoku University.

### Selection and analysis of donor individuals

The details of participant selection were described in our previous study (8). Briefly, three adult male Japanese volunteers were recruited. Japanese ancestry confirmation and individuals self-reported being healthy without any genetic diseases were obtained. Principal Component Analysis (PCA), which verifies their similarity with the Japanese population, and G-band analysis, which validates normal karyotypes, were shown in the previous study (8).

### PacBio CLR

The details of PacBio CLR sequencing were described previously (8). Genomic DNA extracted from nucleated blood cells was fragmented to approximately 20 kb in size and utilized for library preparation using a DNA template prep kit 2.0 from Pacific Biosciences (Menlo Park, CA). Size selection was performed using the Blue Pippin system (Sage Science; Beverly, MA), targeting fragments of 18 kb (with exceptions of 10–15 kb for certain libraries of jg1a). The libraries were then sequenced on a PacBio RSII instrument utilizing P6-C4 chemistry.

### Bionano optical genome mapping

Optical genome mapping was carried out using either the Irys system or the Saphyr system, following the manufacturer’s protocol provided by Bionano Genomics (San Diego, CA). The details of Bionano optical genome mapping were also described in the previous study (8).

### Mate-pair dataset

Genomic DNA extracted from nucleated blood cells was utilized for library construction using a Nextera Mate Pair Library Preparation kit from Illumina, following the gel-free protocol provided by the manufacturer. This protocol yields a broader range of fragment sizes from 2 to 15 kb. Subsequently, the obtained libraries underwent size selection to achieve a range of 300–800 bp (with a peak at 500 bp) using AMPure XP beads from Beckman Coulter (Indianapolis, IN). The libraries were then sequenced on a HiSeq 2500 system from Illumina, employing a TruSeq Rapid PE Cluster kit and TruSeq Rapid SBS kit to obtain 201-bp paired-end reads. Mate-pair dataset 1 and dataset 2 were sequenced under the sequencing depth of 12-13 and 34-38, respectively.

### Short-read paired-end

Short-read paired-end sequencing methods are consistent with those described in previous research (8). Specifically, genomic DNA extracted from buffy coat samples was fragmented to an average target size of 550 bp. Library construction was then performed using the TruSeq DNA PCR-Free HT sample prep kit (Illumina, San Diego, CA), followed by sequencing on a HiSeq 2500 system. This utilized the TruSeq Rapid PE Cluster kit and TruSeq Rapid SBS kit to generate 162- or 259-bp paired-end reads.

### Oxford Nanopore Technologies long reads

Details of sample preparation and sequencing method of Oxford Nanopore Technologies were described previously (8). Genomic DNA was extracted from whole blood using the Gentra Puregene Blood kit (Qiagen). Libraries were prepared with the SQK-LSK109 ligation kit (Oxford Nanopore Technologies) and sequenced on MinION devices using R9.4.1 flow cells. Base-calling was performed with Guppy software (v3.2.4), retaining reads with a Phred quality score above 6 after cropping 100 bp from both ends for mapping.

### Hi-C

Hi-C experiments were essentially performed according to a previously published protocol (15). In brief, five million cells were cross-linked with 1% formaldehyde and quenched with 0.2 M glycine. Cells were lysed using Hi-C lysis buffer (10 mM Tris-HCl pH 8.0, 10 mM NaCl, 0.2% Igepal CA-630), and the chromatin was digested either by MboI (NEB, R0147) or HindIII-HF (NEB). Both ends of the digested chromatin were filled in and labeled with biotin-14-dATP (Life Technologies) for MboI or biotin-14-dCTP (Life Technologies) for HindIII-HF using Klenow Fragment (NEB) and ligated with T4 DNA Ligase (NEB). The biotin-labeled DNA was treated with Proteinase K, reverse cross-linked, and sheared to 300– 500 bp using a Covaris S220 Focused-ultrasonicator. After the size selection using AMPure XP beads (Beckman Coulter), biotin-labeled sheared DNA fragments were enriched using Dynabeads MyOne Streptavidin T1 beads (Life Technologies). The recovered DNA was end-repaired and ligated to Illumina indexed adapters using the NEBNext Ultra DNA Library Prep Kit for Illumina NEB) and NEBNext Multiplex Oligos for Illumina (Index Primers Set 1: NEB). The adapter-ligated DNA underwent 6 or 8 cycles of PCR amplification, followed by AMPure XP bead purification, and then used for sequencing.

### De novo assembly of PacBio subreads

For each individual, we performed a phased assembly using PacBio CLR reads by FALCON, FALCON-unzip (ver. 1.1.2), and Quiver software of the pb-assembly software suite (14). By using FALCON for initial assembly, primary contigs and associated contigs were generated. The results underwent full diploid assembly by FALCON-unzip. The outputs represented the updated primary contigs and haplotype-specific contigs as haplotigs. Subsequently, the results underwent genomic consensus calling by Quiver, yielded the polished version of primary contigs and haplotigs.

### Phased assembly

To address the problem of haplotype switching, we employed phased assembly by FALCON-Phase (ver. 1.1.0) (16). Partially phased long-read assemblies, composed of primary contigs and haplotigs obtained from FALCON-unzip, along with genome-wide chromatin interaction datasets from Hi-C data, were used as the inputs for the analysis. Briefly, the area where a haplotig intersects a primary contig constitutes a phase block, while sections of the primary contig devoid of associated haplotigs are denoted as collapsed regions. Primary contigs undergo segmentation at the alignment start and end positions of haplotigs. Hi-C read pairs are aligned to these segmented contigs, with only haplotype-specific alignments retained. A phasing algorithm assigns phase blocks to either state 0 or 1. FALCON-Phase generates two complete pseudo-haplotypes representing phases 0 and 1. Since two sets of Hi-C data deriving from MboI and HindIII enzymes were used, four sets of fully PacBio/Hi-C phased contigs were obtained from this step for each individual.

### Scaffolding by SALSA2

SALSA2 (ref.15) (version 2.2), a scaffolding tool utilizing genomic proximity information from Hi-C datasets, was employed for the scaffolding process of fully phased contigs. At this step, the Hi-C dataset that was used for each fully phased contigs is the Hi-C dataset derived from different restriction enzymes from the previous phasing step. In other words, the MboI Hi-C dataset was used for scaffolding the HindIII-based phased assemblies, and conversely, the HindIII Hi-C dataset was used for scaffolding the MboI-based phased assemblies. Finally, four sets of PacBio/Hi-C phased scaffolds were obtained for each individual.

### De novo assembly of Bionano optical genome maps

We obtained two sets of Bionano optical genome maps using two enzymes, Nt.BspQI and Nb.BssSI, for subject jg1a, and one set of Bionano optical genome maps was obtained with DLE-1 for jg1b and jg1c. In both cases, the Bionano optical genome maps were assembled in two steps—a rough assembly step and a full assembly step—to perform de novo assembly as independently as possible from the reference. The BionanoSolve software suite (ver. 3.2.1, ver. 3.5) was used for computation.

### De novo assembly of nanopore reads

*De novo* assembly of Oxford Nanopore Technologies (ONT) nanopore reads was conducted using Shasta (0.3.0), Racon (GitHub commit tag 6ca733a), and Medaka (ver. 0.11.1) software.

### Polishing with Pilon

Two sets of Illumina paired-end short reads, 162 bp and 259 bp, were aligned to the fully phased contigs and hybrid scaffolds utilizing BWA MEM software (17) (version 0.7.17). The resulting alignment files were sorted by coordinates and compressed using the Picard tools - SortSam command (version 2.20.5). Subsequently, the BAM files for the 162- and 259-bp paired-end reads were merged using the Picard tools MergeSamFiles command. These merged BAM files were then split into individual scaffolds using the SAMtools (18) (version 1.9) view command, after which each contig/scaffold underwent polishing using Pilon software (19) (version 1.23). Finally, the polished contig/scaffolds were merged into a single multi-FASTA format file.

### Meta-assembly

The Metassembler algorithm (20) performs pairwise, progressive alignments to merge multiple assemblies in the order specified by the user. One of the input assemblies will be used as the primary assembly and the other as the secondary assembly. Mate-pair sequences dataset 1 and dataset 2 were used for meta-assembly within each individual and between individuals, respectively. The compression–expansion statistic (CE statistic) is calculated in both primary and secondary assemblies based on the mapping of mate-pair sequences in each assembly. The data from the secondary assembly are used to improve the primary assembly, such as correction of insertion/deletion errors, closing gaps, and scaffolding sequences, which are based on comparing CE statistics between two assemblies.

For meta-assembly within an individual, four sets of PacBio/Hi-C phased scaffolds from each individual underwent a meta-assembly process by using the Metassembler software (ver. 1.5 with the modification described in Takayama et al., 2019 (ref.8)) (20). There were 24 possible combinations of meta-assembly of four sets of phased scaffolds (**Supplementary Table 5**). Among 24 meta-assemblies, one meta-assembly with the longest scaffold length and the least number of scaffolds was selected for further hybrid scaffolding with Bionano-assembled genome maps and ONT assembly.

For meta-assembly among the three individuals, the three sets of polished scaffolds were then meta-assembled using Metassembler software (20). There were 12 possible combinations to meta-assemble the three sets: (jg1a + (jg1b + jg1c)), (jg1a + (jg1c + jg1b)), ((jg1a + jg1b) + jg1c), ((jg1a + jg1c) + jg1b), (jg1b + (jg1a + jg1c)), (jg1b + (jg1c + jg1a)), ((jg1b + jg1a) + jg1c), ((jg1b + jg1c) + jg1a), (jg1c + (jg1a + jg1b)), (jg1c + (jg1b + jg1a)), ((jg1c + jg1a) + jg1b), and ((jg1c + jg1b) + jg1a), where x + y indicates meta-assemble x and y in this order. For each round of meta-assembly, assemblies were aligned using NUCmer (21), filtered with delta-filter, and converted to COORDS format using show-coords. Mate-pair reads were classified with NxTrim (22) and mapped using Bowtie2 (ref.23), followed by processing with mateAn. The alignment and mapping information were integrated using asseMerge, and the final output was converted to FASTA format with meta2fasta. All of the meta-assemblies in this step were used for anchoring to generate pseudo-molecules.

### Detection of *in silico* STS marker amplification and anchoring scaffolds to chromosomes

We performed *in silico* amplification of the STS markers of three genetic and six RH maps (Genethon (24), Marshfield (25), and deCODE (26) genetic maps; GeneMap-G3 (ref.27), GeneMap99-GB4 (ref.27), TNG (28), NCBI_RH (29), Stanford-G3 (ref.30), and Whitehead-RH maps (31)) on the meta-scaffolds by using in-house electronic PCR software, gPCR (version 2.6a). The STS markers were sourced from the UniSTS database (ftp://ftp.ncbi.nih.gov/pub/ProbeDB/legacy_unists/). One combination of meta-scaffolds from jg1a, jg1b, and jg1c per each chromosome was selected for downstream analysis. The meta-scaffolds were anchored to the chromosomes by the path command of ALLMAPS (ver. 0.8.12) (32).

### Mitochondrial genome and unplaced scaffolds

For the mitochondrial genome, we utilized the mitochondrial genome from JG1. Unanchored contigs/scaffolds generated from anchoring processes against the genetic and radiation hybrid maps using ALLMAPS were also collected. Subsequently, we removed contigs/scaffolds that mapped to the previously described set of chromosomes. Additionally, contigs/scaffolds shorter than 1 kb were excluded. Next, we conducted an all-by-all alignment of the remaining contigs/scaffolds to obtain a set of unique sequences. Finally, scaffolds with N-gaps exceeding 80% of the sequence were excluded.

### Major allele substitution and manual modification

The selected set of meta-assemblies was aligned with two sets of six meta-scaffolds (HindIII and MBoI) using minimap2 (version 2.17) (33). Variants in each set were identified using the paftools call command. To standardize variant representation, we used the BCFtools norm command (version 1.9). Major allele substitutions were conducted by selecting the allele shared by more than three of six meta-scaffolds in each set. For multi-allelic sites with equal allele frequencies, the selection was made randomly. Consecutive N-gap length for heterochromatic regions was manually modified.

### JG2 assembly assessment Consensus quality

The JG2 was aligned with the GRCh38 reference genome using the NUCmer command from the MUMmer software suite (21). The proportion of covered regions and the average identity between the genomes were calculated using the dnadiff tool, also part of the MUMmer suite (34). Additionally, assemblies were aligned to the GRCh38 reference genome using minimap2 software, and variants were identified with the paftools call command. The normalized variants were then annotated using the SnpEff software (35), referencing the GRCh38.86 database.

### Representativeness of JG2 for Japanese variants

To determine whether JG2 harbors the major allele among the Japanese, JG2 was aligned against the reference genome hs37d5 to detect SNVs. Genome-by-genome alignment and comparison were performed using minimap2 and paftools software (33). Allele frequency (AF) was investigated on the 3.5KJPNv2 AF panel (36, 37). Allele frequency spectrums of JG2 and JG1 were created with the horizontal axis representing the non-hs37d5-type allele and the vertical axis showing the number of such variant sites.

## Results

### JG2 construction

The individuals selected for constructing JG2 were three Japanese male volunteers named jg1a, jg1b, and jg1c, the same individuals used to create the previous Japanese reference genome, JG1 (ref.8). For each individual, we obtained over 120× PacBio CLR, two sets of over 49× Hi-C reads (using MboI and HindIII enzymes), one or two sets of over 120× Bionano optical genome maps (using DSL-1 or BspQI and BssSI enzymes), one set of ONT long reads, two sets of mate-pair read, and two sets of paired-end reads (Illumina HiSeq 162-bp and 259-bp reads). The depth of reads or optical genome maps is summarized in **Supplementary Table 1.**

### Genome assembly for each individual

JG2 was created by first performing a phased assembly for each individual and then integrating genomes among the three Japanese individuals (**Figure 1**). We performed a phased assembly for each individual using PacBio CLR reads by Falcon, Falcon Unzip, and Quiver software of the pb-assembly software suite (**Figure 1A**). The assembly statistics, including total length, N50, and the number of contigs, were in **Supplementary Table 2**. After the polishing, the number of primary contigs of jg1a, jg1b, and jg1c were 1 439, 1 386, and 1 271, respectively. The N50 of all participants is around 20 Mb.

**Figure 1.**
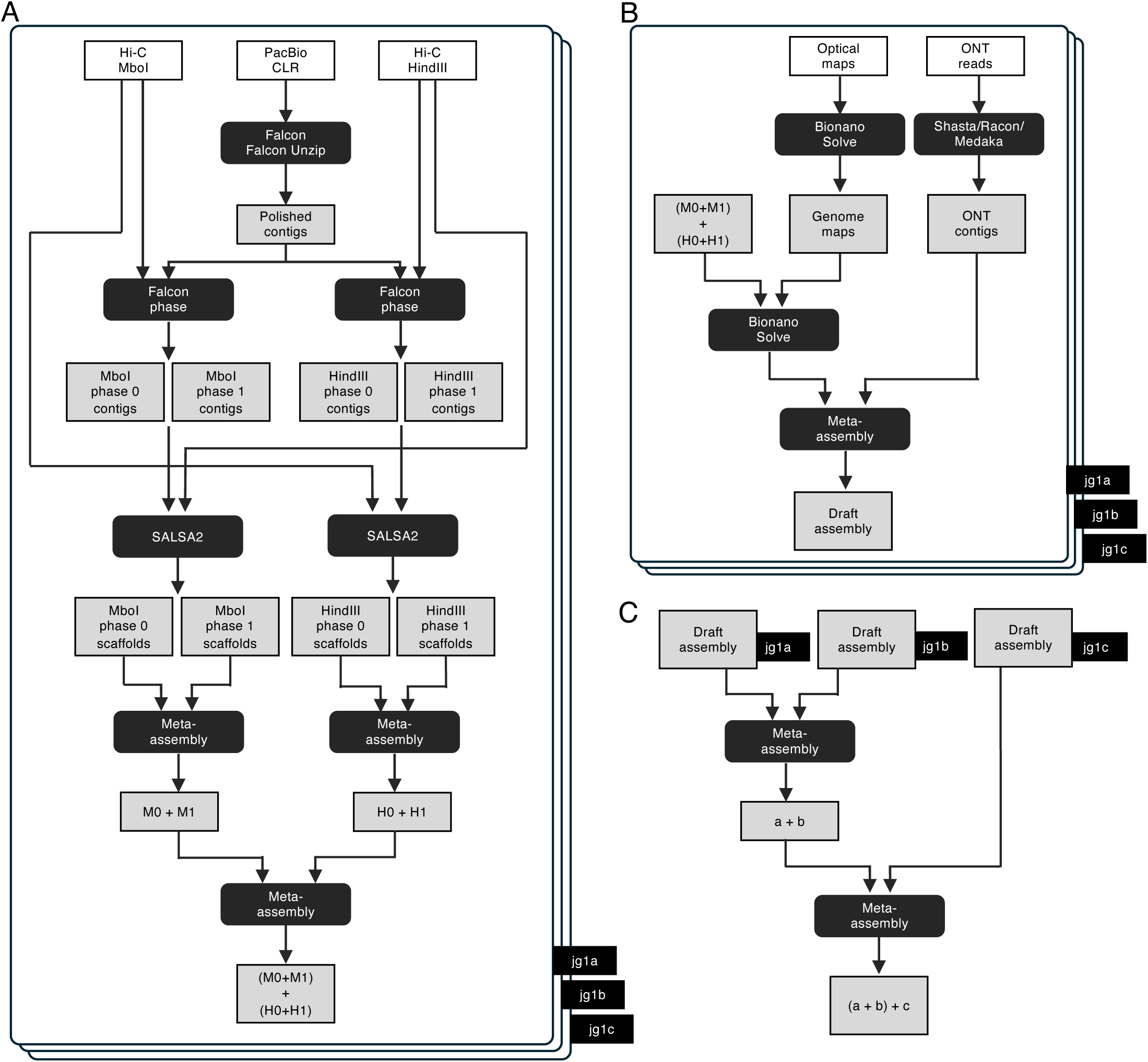
Construction strategy of the JG2 reference genome. Three Japanese male volunteers (jg1a, jg1b, jg1c) provided the data for JG2. High-coverage sequencing and optical genome mapping data were generated: PacBio CLRs, Hi-C reads, Bionano optical genome maps, ONT nanopore long reads, Illumina mate-pair reads, and Illumina paired-end reads. **(A)** Phased genome assemblies and scaffolding were performed for each individual using PacBio CLRs and Hi-C reads. Meta-assemblies were created within each individual. **(B)** Bionano optical genome maps were assembled, and PacBio/Hi-C-based assemblies were scaffolded with the optical genome maps in each individual. In addition, ONT long reads were assembled within each individual, then meta-assembled with the PacBio/Hi-C/Bionano-based assemblies, resulting in a draft assembly for each individual. **(C)** Draft assemblies for the three individuals were meta-assembled. White boxes indicate raw data, gray boxes indicate intermediate assemblies, and black rounded boxes indicate software or operations. CLR, continuous long reads; ONT, Oxford Nanopore Technology.

Subsequently, by using two sets of Hi-C data separately, we conducted phased assembly, which generated two sets of haploid genome assemblies per individual using Falcon Phase software: MboI-based phased assemblies and HindIII-based phased assemblies (**Figure 1A**). The assembly statistics are shown in **Supplementary Table 3**. These assemblies were then polished using 162-bp and 259-bp HiSeq short reads by Pilon software. Scaffolding was conducted on each set of polished haploid genome assemblies of the PacBio/Hi-C phased contigs utilizing SALSA2 software. Specifically, we utilized the Hi-C dataset that was not employed for phasing. For example, we used the MboI Hi-C dataset for scaffolding the HindIII-based phased assemblies. Conversely, we used the HindIII Hi-C dataset for scaffolding the MboI-based phased assemblies (**Figure 1A**). Consequently, we obtained four sets of PacBio/Hi-C phased scaffolds for each individual. The assembly statistics are shown in **Supplementary Table 3**. The number of scaffolds of four sets of phased scaffolds in jg1a, jg1b, and jg1c was 997 ± 55 (mean ± s.d.).

In addition to PacBio/Hi-C phased scaffolds, de novo assembly of Bionano optical genome maps was performed by BionanoSolve software, and de novo assembly of ONT nanopore reads was conducted using Shasta, Racon, and Medaka software (**Figure 1B**). Subsequently, the ONT assemblies were refined by polishing with 162-bp and 259-bp HiSeq short reads using Pilon software. The summary of assembly statistics is presented in **Supplementary Table 4**.

In summary we acquired four sets of phased PacBio/Hi-C assemblies from each participant, along with one or two sets of Bionano-assembled genome maps and one set of ONT assemblies. These datasets were subsequently used in further meta-assembly.

### Meta-assembly within each individual

The meta-assembly process involves integrating the locally best sequences from all input assemblies across the genome. These sequences are merged to create a final sequence that is either as good as or superior to the individual constituent assemblies. For each individual, four sets of phased PacBio/Hi-C assemblies underwent two-step meta-assemblies, resulting in 24 different meta-assemblies (**Figure 1A**). Then, one meta-assembly that had the least number of mis-assemblies when compared with genetic/RH maps and JG1 was selected for further hybrid scaffolding with Bionano assembled genome maps and ONT assembly. Mate-pair short reads were used for meta-assembly by Metassembler software. The meta-assembly statistics for two-step meta-assemblies of phased PacBio/Hi-C assemblies are shown in **Supplementary Table 5**.

Following that, we conducted hybrid scaffolding of the meta-assembly using Bionano genome maps through BionanoSolve software. Subsequently, the resulting hybrid scaffolds were merged with the ONT assembly utilizing merged dataset 1 mate-pair short reads via Metassembler software. In this step of meta-assemblies within each individual, one meta-assembly per individual was generated. Consequently, we obtained three sets of meta-assemblies for the three individuals.

### Meta-assemblies among individuals

Meta-assembly was conducted among the three individuals using mate-pair short reads from dataset 2 through Metassembler software, resulting in 12 sets of meta-assemblies (**Figure 1C**). Subsequently, 12 sets of meta-assemblies were anchored to 3 genetic and 6 radiation hybrid maps using ALLMAPS software, resulting in 12 sets of pseudo-molecules.

### Selection of JG2

By comparing the pseudo-molecules anchored to each linkage group to the corresponding chromosome sequence of JG2, one pseudo-molecule was chosen for each chromosome sequence for JG2 (**Supplementary Table 6**). The selected pseudo-molecules, mitochondrial reference genome, and unplaced scaffolds comprise a full set of the reference genome. Major allele substitution was performed using variants from two sets of six meta-scaffolds (HindIII and MBoI) to remove rare individual variants. Manual modifications were conducted to adjust the N-gap length for the acrocentric, centromeric, heterochromatic, and telomeric regions, yielding the final set of genome sequences of JG2.

### Evaluation of JG2

The procedure described above resulted in a set of chromosome-level sequences for 22 autosomes, 2 sex chromosomes, 1 mitochondrial chromosome, and 1148 unplaced scaffolds collectively designated as JG2. The total length of JG2 was approximately 3.1 Gb, which included 609 gap regions totaling 251 Mb. Of these gaps, 233.63 Mb were intentionally inserted to represent telomeric, centromeric, and heterochromatic regions. The scaffold N50 was 152 668 378 bp. The number of misassemblies was 1 055, respectively. Alignments between JG2 and several reference genomes, including GRCh38, hs37d5, and the previously constructed JG1, are shown in **Figure 2**. JG2 covered 95.35% of the reference genome with an average identity of 99.80%.

**Figure 2.**
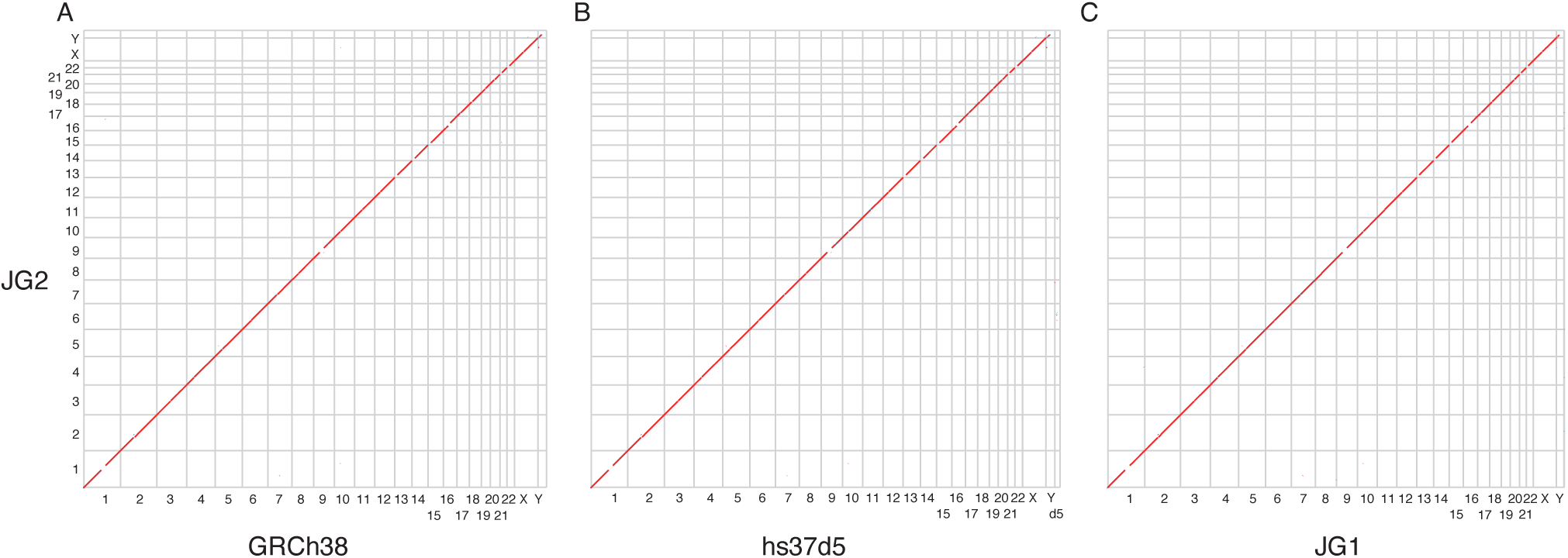
Dot plots show co-linearity between JG2 and other reference genomes. **(A)** JG2 vs. GRCh38. **(B)** JG2 vs. hs37d5. **(C)** JG2 vs. JG1.

When compared to the GRCh38 reference genome, JG2 contained 3 115 695 variants, of which 2 130 (0.054%) were classified as HIGH-impact variants. The number of protein-truncating variants (PTVs), which include stop-gained, frameshift, splice-acceptor, and splice-donor variants, were 55, 280, 141, and 118, respectively.

### Representativeness of JG2 for SNVs composition of the Japanese population

The genome-by-genome alignment and comparison revealed 2 321 710 SNVs between hs37d5 and JG2 in the autosomes and X chromosome. Of these SNVs, 298 644 had an allele frequency (AF) of 1.0 in the 3.5KJPNv2 AF panel, indicating that all Japanese have JG2-type alleles at these 298 644 sites. This number is larger than that of JG1 (246 464), demonstrating that JG2 better represents major alleles in the Japanese population. Additionally, we identified 366 364 SNV sites with an AF ≥ 0.99 and 624 847 SNV sites with an AF ≥ 0.90 in the 3.5KJPNv2 AF panel, respectively. Allele frequency spectrums for SNVs comparing JG2 and JG1 are demonstrated in **Figure 3**.

**Figure 3.**
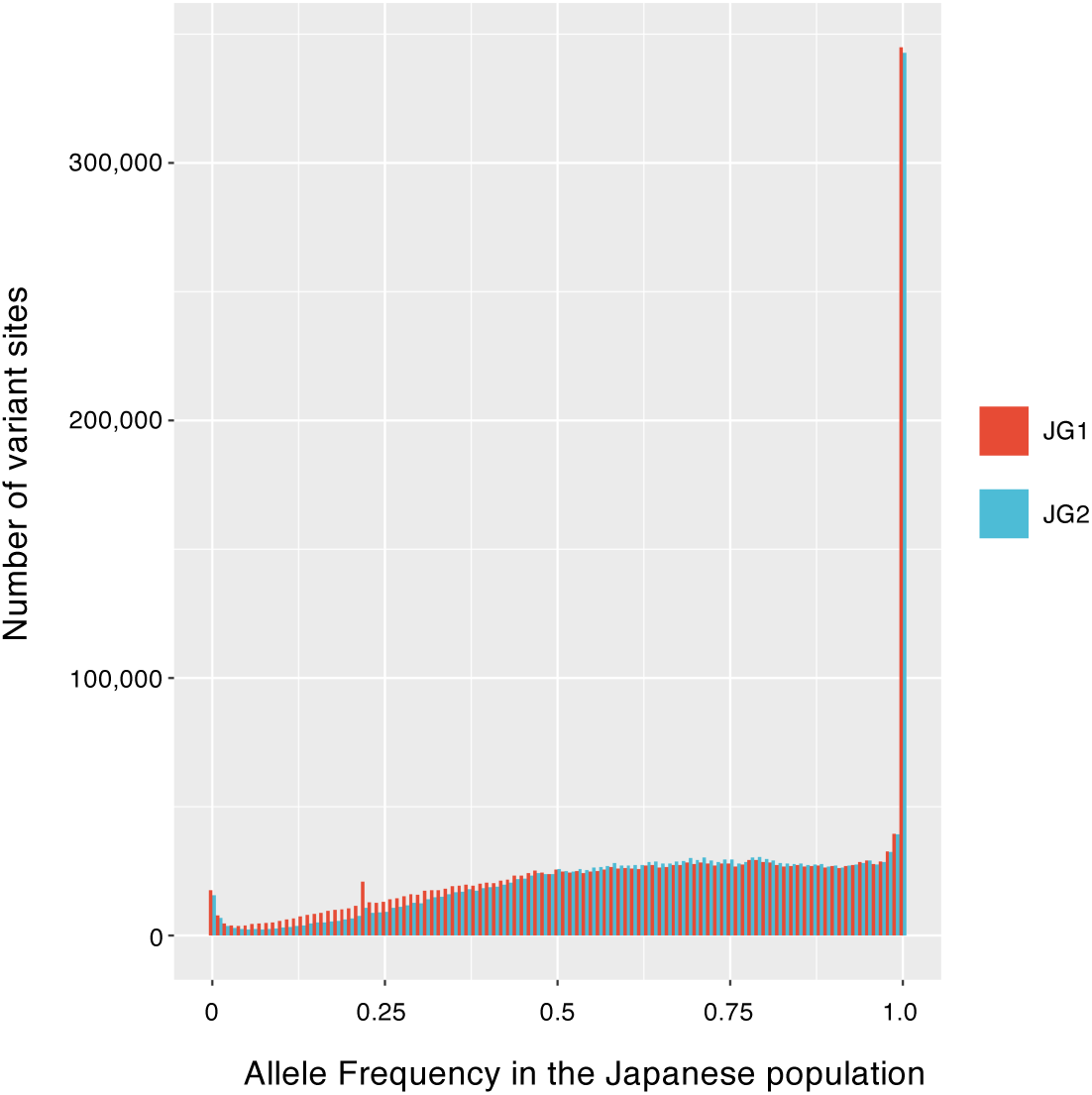
The allele frequency spectrum of JG2 and JG1. JG2 was aligned to the reference genome hs37d5 to detect SNVs. Allele frequencies in the Japanese population were analyzed using the 3.5KJPNv2 allele frequency panel. The spectrum displays the alternative allele frequency for SNVs on the horizontal axis and the number of variant sites on the vertical axis.

## Discussion

The current study demonstrated the strategy to assemble the population-specific reference genome, JG2, from three individuals whose genome data were used to construct the previous version of the Japanese specific reference genome, JG1. By applying advanced strategies, JG2 better covers Japanese-specific variants and enhances assembly statistics. Using Hi-C data for phase-aware assembly, the process generated two haploid assemblies per individual. Unlike JG1, this method treats the haploid genomes as random samples from a population, increasing genetic variation representation. The increase in haplotypes enhanced population representativeness and, through majority decision, reduced the presence of rare personal variants, making JG2 more generalizable to the Japanese population.

Hi-C sequencing technology was instrumental in our phased assembly process. Hi-C data provides long-range contact information that helps to accurately place contigs and scaffolds into their correct positions along the chromosomes, enabling the differentiation of maternal and paternal haplotypes (38, 39). This phasing is crucial for understanding the allelic variations and structural differences between the two sets of chromosomes, providing a more complete and accurate representation of the genome.

The meta-assembly strategy, which involved combining the best sequences from multiple assemblies, enhanced assembly quality for the Japanese genome (8). For each individual, the meta-assembly process led to enhanced scaffold lengths, indicated by the increased N50 in JG2 compared to JG1 (152 668 378 vs 141 953 703). In addition, a majority decision strategy was also applied to remove rare variants. By comparing multiple assemblies and subsequent selection of the consensus sequence, we effectively mitigated the inclusion of sequencing errors and rare personal variants that might not represent the population. This approach reduced the number of misassemblies in JG2 compared to JG1 (1 055 vs 1 654). This approach ensured that the final assemblies were more accurate and reflective of the common genomic features within the Japanese population. The iterative meta-assembly among individuals further consolidated these improvements, creating comprehensive and representative pseudo-molecules for each chromosome.

Despite its advancements, JG2 is based on technology slightly behind the state-of-the-art compared to the recent developments presented by T2T (40) and pangenome projects (41, 42). T2T utilized high-accuracy long-read sequences such as HiFi reads to construct a high-resolution assembly string graph to address repetitive and complex regions such as centromeres, telomeres, and other difficult-to-sequence areas (40). A pangenome encompasses the collective whole-genome sequences of multiple individuals to capture the genetic diversity of a species, utilizing high-quality, phased haplotypes and a graph-based data structure for improved reference and variant indexing, with compatibility to existing human reference genomes (41, 42). However, it is important to note that JG2 has been steadily updated compared to JG1. This continuous improvement ensures that JG2 remains a valuable fundamental information resource that can be used for various analyses, such as identifying population-specific genetic variations or conducting comparative studies within the Japanese population.

In conclusion, this study improved the Japanese reference genome, creating JG2 via phased assembly from three individuals. Hi-C data enabled phase-aware assembly, generating two haploid assemblies per individual and better representing genetic variation. A meta-assembly strategy combined the best sequences, improving scaffold lengths and accuracy, while a majority decision approach minimized rare variants. These methods produced high-quality, comprehensive pseudo-molecules, making JG2 more representative of the Japanese population.

## Data availability

The JG2 sequences, along with additional resources, are available from the jMorp website (37) (https://jmorp.megabank.tohoku.ac.jp/downloads/tommo-jg2.0.0.beta-20200831).

## Acknowledgments

This work was supported in part by the Tohoku Medical Megabank (TMM) Project from the Ministry of Education, Culture, Sports, Science and Technology (MEXT) and by the Japan Agency for Medical Research and Development (AMED; Grant Numbers JP21tm0124005) for Tohoku University. This work was also supported in part by JST Moonshot R&D Program Grant Number JPMJMS2023 to GT and JT. All computational resources were provided by the ToMMo supercomputer system (http://sc.megabank.tohoku.ac.jp/en), which is supported by Facilitation of R&D Platform for AMED Genome Medicine Support conducted by AMED (Grant Number JP21tm0424601). We appreciate all the volunteers who participated in the TMM project.

## Supplementary information

Supplementary information is available at Journal of Human Genetics’ website

## References

1. Lander ES, Linton LM, Birren B, Nusbaum C, Zody MC, Baldwin J, et al. Initial sequencing and analysis of the human genome. Nature. 2001;409(6822):860–921.

2. Venter JC, Adams MD, Myers EW, Li PW, Mural RJ, Sutton GG, et al. The sequence of the human genome. Science. 2001;291(5507):1304–51.

3. Ameur A, Che H, Martin M, Bunikis I, Dahlberg J, Höijer I, et al. De Novo Assembly of Two Swedish Genomes Reveals Missing Segments from the Human GRCh38 Reference and Improves Variant Calling of Population-Scale Sequencing Data. Genes (Basel). 2018;9(10).

4. Cho YS, Kim H, Kim H-M, Jho S, Jun J, Lee YJ, et al. An ethnically relevant consensus Korean reference genome is a step towards personal reference genomes. Nature Communications. 2016;7(1):13637.

5. Seo J-S, Rhie A, Kim J, Lee S, Sohn M-H, Kim C-U, et al. De novo assembly and phasing of a Korean human genome. Nature. 2016;538(7624):243–7.

6. Shi L, Guo Y, Dong C, Huddleston J, Yang H, Han X, et al. Long-read sequencing and de novo assembly of a Chinese genome. Nature Communications. 2016;7(1):12065.

7. Ouzhuluobu, He Y, Lou H, Cui C, Deng L, Gao Y, et al. De novo assembly of a Tibetan genome and identification of novel structural variants associated with high-altitude adaptation. National Science Review. 2019;7(2):391–402.

8. Takayama J, Tadaka S, Yano K, Katsuoka F, Gocho C, Funayama T, et al. Construction and integration of three de novo Japanese human genome assemblies toward a population-specific reference. Nature Communications. 2021;12(1):226.

9. Uneoka S, Kobayashi T, Numata-Uematsu Y, Oikawa Y, Katata Y, Okubo Y, et al. A Case Series of Patients With MYBPC1 Gene Variants Featuring Undulating Tongue Movements as Myogenic Tremor. Pediatric Neurology. 2023;146:16–20.

10. Shibuya M, Yaoita H, Kodama K, Okubo Y, Endo W, Inui T, et al. A patient with early-onset SMAX3 and a novel variant of ATP7A. Brain and Development. 2022;44(1):63–7.

11. Katata Y, Uneoka S, Saijyo N, Aihara Y, Miyazoe T, Koyamaishi S, et al. The longest reported sibling survivors of a severe form of congenital myasthenic syndrome with the pathogenic variant. American Journal of Medical Genetics Part A. 2022;188(4):1293–8.

12. Ito S, Hashimoto A, Yamaguchi K, Kawamura S, Myoen S, Ogawa M, et al. A novel 8.57-kb deletion of the upstream region of PRKAR1A in a family with Carney complex. Molecular Genetics & Genomic Medicine. 2022;10(3):e1884.

13. Chin C-S, Peluso P, Sedlazeck FJ, Nattestad M, Concepcion GT, Clum A, et al. Phased diploid genome assembly with single-molecule real-time sequencing. Nature Methods. 2016;13(12):1050–4.

14. Kronenberg ZN, Rhie A, Koren S, Concepcion GT, Peluso P, Munson KM, et al. Extended haplotype-phasing of long-read de novo genome assemblies using Hi-C. Nature Communications. 2021;12(1):1935.

15. Ghurye J, Rhie A, Walenz BP, Schmitt A, Selvaraj S, Pop M, et al. Integrating Hi-C links with assembly graphs for chromosome-scale assembly. PLOS Computational Biology. 2019;15(8):e1007273.

16. Lieberman-Aiden E, van Berkum NL, Williams L, Imakaev M, Ragoczy T, Telling A, et al. Comprehensive mapping of long-range interactions reveals folding principles of the human genome. Science. 2009;326(5950):289–93.

17. Li H. Aligning sequence reads, clone sequences and assembly contigs with BWA-MEM. arXiv: Genomics. 2013.

18. Li H, Handsaker B, Wysoker A, Fennell T, Ruan J, Homer N, et al. The Sequence Alignment/Map format and SAMtools. Bioinformatics. 2009;25(16):2078–9.

19. Walker BJ, Abeel T, Shea T, Priest M, Abouelliel A, Sakthikumar S, et al. Pilon: An Integrated Tool for Comprehensive Microbial Variant Detection and Genome Assembly Improvement. PLOS ONE. 2014;9(11):e112963.

20. Wences AH, Schatz MC. Metassembler: merging and optimizing de novo genome assemblies. Genome Biology. 2015;16(1):207.

21. Marçais G, Delcher AL, Phillippy AM, Coston R, Salzberg SL, Zimin A. MUMmer4: A fast and versatile genome alignment system. PLOS Computational Biology. 2018;14(1):e1005944.

22. O’Connell J, Schulz-Trieglaff O, Carlson E, Hims MM, Gormley NA, Cox AJ. NxTrim: optimized trimming of Illumina mate pair reads. Bioinformatics. 2015;31(12):2035–7.

23. Langmead B, Salzberg SL. Fast gapped-read alignment with Bowtie 2. Nature Methods. 2012;9(4):357–9.

24. Dib C, Fauré S, Fizames C, Samson D, Drouot N, Vignal A, et al. A comprehensive genetic map of the human genome based on 5,264 microsatellites. Nature. 1996;380(6570):152–4.

25. Broman KW, Murray JC, Sheffield VC, White RL, Weber JL. Comprehensive human genetic maps: individual and sex-specific variation in recombination. Am J Hum Genet. 1998;63(3):861–9.

26. Kong A, Gudbjartsson DF, Sainz J, Jonsdottir GM, Gudjonsson SA, Richardsson B, et al. A high-resolution recombination map of the human genome. Nature Genetics. 2002;31(3):241–7.

27. Stewart EA, McKusick KB, Aggarwal A, Bajorek E, Brady S, Chu A, et al. An STS-based radiation hybrid map of the human genome. Genome Res. 1997;7(5):422–33.

28. Olivier M, Aggarwal A, Allen J, Almendras AA, Bajorek ES, Beasley EM, et al. A high-resolution radiation hybrid map of the human genome draft sequence. Science. 2001;291(5507):1298–302.

29. Agarwala R, Applegate DL, Maglott D, Schuler GD, Schäffer AA. A fast and scalable radiation hybrid map construction and integration strategy. Genome Res. 2000;10(3):350–64.

30. Deloukas P, Schuler GD, Gyapay G, Beasley EM, Soderlund C, Rodriguez-Tomé P, et al. A physical map of 30,000 human genes. Science. 1998;282(5389):744–6.

31. Hudson TJ, Stein LD, Gerety SS, Ma J, Castle AB, Silva J, et al. An STS-based map of the human genome. Science. 1995;270(5244):1945–54.

32. Tang H, Zhang X, Miao C, Zhang J, Ming R, Schnable JC, et al. ALLMAPS: robust scaffold ordering based on multiple maps. Genome Biology. 2015;16(1):3.

33. Li H. Minimap2: pairwise alignment for nucleotide sequences. Bioinformatics. 2018;34(18):3094–100.

34. Phillippy AM, Schatz MC, Pop M. Genome assembly forensics: finding the elusive mis-assembly. Genome Biology. 2008;9(3):R55.

35. Cingolani P, Platts A, Wang le L, Coon M, Nguyen T, Wang L, et al. A program for annotating and predicting the effects of single nucleotide polymorphisms, SnpEff: SNPs in the genome of Drosophila melanogaster strain w1118; iso-2; iso-3. Fly (Austin). 2012;6(2):80–92.

36. Tadaka S, Katsuoka F, Ueki M, Kojima K, Makino S, Saito S, et al. 3.5KJPNv2: an allele frequency panel of 3552 Japanese individuals including the X chromosome. Human Genome Variation. 2019;6(1):28.

37. Tadaka S, Kawashima J, Hishinuma E, Saito S, Okamura Y, Otsuki A, et al. jMorp: Japanese Multi-Omics Reference Panel update report 2023, Nucleic Acids Research. 2024;52(D1):D622–D632.

38. Rao SS, Huntley MH, Durand NC, Stamenova EK, Bochkov ID, Robinson JT, et al. A 3D map of the human genome at kilobase resolution reveals principles of chromatin looping. Cell. 2014;159(7):1665–80.

39. Dudchenko O, Batra SS, Omer AD, Nyquist SK, Hoeger M, Durand NC, et al. De novo assembly of the *Aedes aegypti* genome using Hi-C yields chromosome-length scaffolds. Science. 2017;356(6333):92–5.

40. Nurk S, Koren S, Rhie A, Rautiainen M, Bzikadze AV, Mikheenko A, et al. The complete sequence of a human genome. Science. 2022;376(6588):44–53.

41. Wang T, Antonacci-Fulton L, Howe K, Lawson HA, Lucas JK, Phillippy AM, et al. The Human Pangenome Project: a global resource to map genomic diversity. Nature. 2022;604(7906):437–46.

42. Liao W-W, Asri M, Ebler J, Doerr D, Haukness M, Hickey G, et al. A draft human pangenome reference. Nature. 2023;617(7960):312–24.

