## Supplementary Information for "JG2: an updated version of the Japanese population-specific reference genome"

**Supplementary Table 1.** Sequencing platform and sequencing depth using in this study

| **Platform** | **jg1a** | **jg1b** | **jg1c** | **Reference** |
| --- | --- | --- | --- | --- |
| PacBio CLR | 122× | 123× | 128× | (8) |
| Bionano | 123× (BspQI)  140× (BssSI) | 160× (DLE-1) | 175× (DLE-1) | (8) |
| Hi-C | 100.3× (MboI)  49.5× (HindIII) | 70.1× (MboI)  50.3× (HindIII) | 71.3× (MboI)  49.2× (HindIII) | This study |
| ONT | 46.2× | 52.0× | 43.9× | (8) |
| mate-pair | 13× (dataset1)  36× (dataset2) | 12× (dataset 1)  38× (dataset 2) | 12× (dataset 1)  34× (dataset 2) | (8) This study |
| paired-end | 29× (162 bp)  26× (259 bp) | 31× (162 bp)  28× (259 bp) | 31× (162 bp)  26× (259 bp) | (8) |

Depth is calculated by assuming the genome size = 3.0 Gb. CLR, continuous long reads; ONT, Oxford Nanopore Technology.

**Supplementary Table 2.** Assembly statistics in initial assembly, unzipping, and polishing steps

| **sample** | **contig** | **assembly**  **(2-asm-falcon)** | | | **unzipping**  **(3-unzip)** | | | **polishing**  **(4-quiver)** | | |
| --- | --- | --- | --- | --- | --- | --- | --- | --- | --- | --- |
|  |  | **Total length** | **N50** | **Number of contigs** | **Total length** | **N50** | **Number of contigs** | **Total length** | **N50** | **Number of contigs** |
| **jg1a** | primary | 2,872,229,904 | 19,759,866 | 1,719 | 2,865,844,877 | 19,764,383 | 1,570 | 2,867,633,543 | 19,806,459 | 1,439 |
|  | associated/haplotig | 126,953,103 | 41,366 | 3,096 | 1,899,278,335 | 235,067 | 15,057 | 1,861,440,548 | 241,839 | 13,015 |
| **jg1b** | primary | 2,866,558,456 | 21,053,413 | 1,663 | 2,860,907,570 | 21,047,859 | 1,526 | 2,862,109,924 | 21,088,092 | 1,386 |
|  | associated/haplotig | 111,302,058 | 36,052 | 3,063 | 1,847,276,479 | 220,720 | 15,203 | 1,817,898,237 | 226,102 | 13,460 |
| **jg1c** | primary | 2,863,290,164 | 18,455,999 | 1,508 | 2,857,954,005 | 18,459,607 | 1,390 | 2,859,587,410 | 18,498,210 | 1,271 |
|  | associated/haplotig | 128,014,679 | 38,090 | 3,405 | 1,909,671,674 | 230,691 | 15,175 | 1,877,734,437 | 235,552 | 13,289 |

**Supplementary Table 3.** Assembly statistics in phase assembly, scaffolding steps

| **sample** | **Set** | **phase** | **Phasing (5-phase)** | | | **Scaffolding (SALSA2)** | | | |
| --- | --- | --- | --- | --- | --- | --- | --- | --- | --- |
|  |  |  | **Total length** | **N50** | **Number of contigs** | **Total length** | **N50** | **Number of contigs** | **Number of mis-assemblies** |
| jg1a | H→M | 0 | 2,868,973,897 | 19,816,507 | 1,423 | 2,868,216,930 | 75,737,371 | 1,058 | 2,209 |
|  |  | 1 | 2,868,386,599 | 19,818,126 | 1,423 | 2,868,515,945 | 57,410,013 | 1,060 | 2,180 |
|  | M→H | 0 | 2,867,671,190 | 19,827,283 | 1,423 | 2,867,793,358 | 62,217,234 | 1,039 | 2,211 |
|  |  | 1 | 2,868,789,306 | 19,807,350 | 1,423 | 2,868,944,936 | 76,103,143 | 1,055 | 2,225 |
| jg1b | H→M | 0 | 2,862,734,228 | 21,095,395 | 1,366 | 2,862,726,396 | 61,249,395 | 1,013 | 2,151 |
|  |  | 1 | 2,862,897,069 | 21,084,959 | 1,366 | 2,862,909,839 | 63,068,230 | 1,010 | 2,216 |
|  | M→H | 0 | 2,862,810,851 | 21,093,630 | 1,366 | 2,862,838,203 | 59,385,241 | 1,001 | 2,187 |
|  |  | 1 | 2,862,820,446 | 21,086,724 | 1,366 | 2,862,836,507 | 63,088,539 | 1,014 | 2,168 |
| jg1c | H→M | 0 | 2,860,063,589 | 18,495,858 | 1,257 | 2,860,083,531 | 62,894,796 | 926 | 2,158 |
|  |  | 1 | 2,860,436,381 | 18,494,518 | 1,257 | 2,860,416,655 | 81,779,830 | 931 | 2,118 |
|  | M→H | 0 | 2,860,603,511 | 18,496,741 | 1,257 | 2,860,619,566 | 62,940,103 | 942 | 2,141 |
|  |  | 1 | 2,859,896,459 | 18,493,635 | 1,257 | 2,859,882,019 | 86,925,497 | 912 | 2,167 |

H→M: Phasing with HindIII Hi-C datasets, then scaffolding with MboI Hi-C datasets; M→H: Phasing with MboI Hi-C datasets, then scaffolding with HindIII Hi-C datasets

**Supplementary Table 4**. Assembly statistics for ONT nanopore reads

| **sample** | **assembly stages** | **number of contigs** | **total length** | **NGA50** | **number of misassemblies** |
| --- | --- | --- | --- | --- | --- |
| **jg1a** | assembly (Shasta) | 4,550 | 2,834,936,073 | 10,175,913 | 1,100 |
|  | 1st polishing (Racon) | 4,550 | 2,828,673,945 | 10,241,589 | 874 |
|  | 2nd polishing (Medaka) | 2,876 | 2,827,100,153 | 10,195,383 | 925 |
| **jg1b** | assembly (Shasta) | 7,904 | 2,839,782,966 | 9,553,835 | 1,109 |
|  | 1st polishing (Racon) | 7,904 | 2,833,862,262 | 10,218,051 | 894 |
|  | 2nd polishing (Medaka) | 3,291 | 2,828,695,661 | 9,904,179 | 893 |
| **jg1c** | assembly (Shasta) | 6,174 | 2,832,610,762 | 9,249,159 | 897 |
|  | 1st polishing (Racon) | 6,174 | 2,826,953,599 | 9,580,139 | 859 |
|  | 2nd polishing (Medaka) | 2,801 | 2,824,650,793 | 9,444,579 | 818 |

**Supplementary Table 5.** Within individual meta-assembly statistics

|  | **jg1a** | | | | **jg1b** | | | | **jg1c** | | | |
| --- | --- | --- | --- | --- | --- | --- | --- | --- | --- | --- | --- | --- |
| **assembly** | **Misassemblies vs** | | **total length** | **N50** | **Misassemblies vs** | | **total length** | **N50** | **Misassemblies vs** | | **total length** | **N50** |
|  | **genetic/RH maps** | **JG1** |  |  | **genetic/RH maps** | **JG1** |  |  | **genetic/RH maps** | **JG1** |  |  |
| (H.0 + H.1) + (M.0 + M.1) | 16 | 4085 | 2867177037 | 75735957 | 24 | 4059 | 2861540671 | 61247162 | 14 | 2816 | 2858610684 | 62893376 |
| (H.0 + H.1) + (M.1 + M.0) | 16 | 4072 | 2867150887 | 75735957 | 20 | 4048 | 2861619372 | 61247162 | 14 | 2798 | 2858838197 | 62893376 |
| (H.0 + M.0) + (H.1 + M.1) | 16 | 4081 | 2867402314 | 75735957 | 20 | 4061 | 2861683747 | 61247162 | 14 | 2816 | 2858730520 | 62893376 |
| (H.0 + M.0) + (M.1 + H.1) | 16 | 4078 | 2867463636 | 75735957 | 24 | 4064 | 2861704097 | 61247162 | 14 | 2809 | 2858973017 | 62893376 |
| (H.0 + M.1) + (H.1 + M.0) | 16 | 4059 | 2867248727 | 75735957 | 20 | 4035 | 2861529553 | 61247162 | 14 | 2811 | 2858868283 | 62893376 |
| (H.0 + M.1) + (M.0 + H.1) | 16 | 4079 | 2867417997 | 75735957 | 24 | 4036 | 2861554963 | 61247162 | 14 | 2804 | 2858771765 | 62893376 |
| (H.1 + H.0) + (M.0 + M.1) | 15 | 4067 | 2867279847 | 57408505 | 31 | 3986 | 2861885057 | 63067780 | 17 | 2768 | 2859023915 | 81778283 |
| (H.1 + H.0) + (M.1 + M.0) | 15 | 4069 | 2867416857 | 57408505 | 31 | 4004 | 2862124168 | 63067780 | 17 | 2773 | 2859286998 | 81778283 |
| (H.1 + M.0) + (H.0 + M.1) | 15 | 4065 | 2867447452 | 57408505 | 31 | 3979 | 2861902975 | 63067780 | 17 | 2765 | 2859162550 | 81778283 |
| (H.1 + M.0) + (M.1 + H.0) | 15 | 4072 | 2867671047 | 57408505 | 31 | 3994 | 2862009415 | 63067780 | 17 | 2769 | 2859276824 | 81778283 |
| (H.1 + M.1) + (H.0 + M.0) | 15 | 4067 | 2867570550 | 57408505 | 31 | 4002 | 2862074798 | 63067780 | 17 | 2772 | 2859253014 | 81778283 |
| (H.1 + M.1) + (M.0 + H.0) | 15 | 4078 | 2867595906 | 57408505 | 31 | 3994 | 2861974949 | 63067780 | 17 | 2778 | 2859440475 | 81778283 |
| (M.0 + H.0) + (H.1 + M.1) | 14 | 4112 | 2867094533 | 62215201 | 23 | 4072 | 2861902602 | 59379041 | 17 | 2820 | 2859713533 | 62937813 |
| (M.0 + H.0) + (M.1 + H.1) | 14 | 4096 | 2866862889 | 62215201 | 23 | 4068 | 2861867946 | 59379041 | 17 | 2823 | 2859504686 | 62937813 |
| (M.0 + H.1) + (H.0 + M.1) | 14 | 4099 | 2866992474 | 62215201 | 23 | 4068 | 2861802406 | 59379041 | 17 | 2822 | 2859404386 | 62937813 |
| (M.0 + H.1) + (M.1 + H.0) | 14 | 4086 | 2866840705 | 62215201 | 23 | 4062 | 2861823335 | 59379041 | 26 | 2816 | 2859464851 | 62937813 |
| (M.0 + M.1) + (H.0 + H.1) | 14 | 4096 | 2866789506 | 62215201 | 23 | 4064 | 2861818212 | 59379041 | 17 | 2812 | 2859309823 | 62937813 |
| (M.0 + M.1) + (H.1 + H.0) | 14 | 4089 | 2866651407 | 62215201 | 23 | 4055 | 2861769305 | 59379041 | 17 | 2819 | 2859563681 | 62937813 |
| (M.1 + H.0) + (H.1 + M.0) | 16 | 4119 | 2868019628 | 76102530 | 30 | 4031 | 2861852855 | 63087330 | 25 | 2779 | 2858716510 | 87660285 |
| (M.1 + H.0) + (M.0 + H.1) | 16 | 4114 | 2868056065 | 76102530 | 30 | 4019 | 2861841829 | 63087330 | 25 | 2783 | 2858565596 | 87638277 |
| (M.1 + H.1) + (H.0 + M.0) | 16 | 4132 | 2868023531 | 76102106 | 30 | 4024 | 2862027763 | 63087330 | 25 | 2789 | 2858854529 | 87660623 |
| (M.1 + H.1) + (M.0 + H.0) | 16 | 4127 | 2868036928 | 76102118 | 30 | 4042 | 2861993783 | 63087330 | 16 | 2794 | 2858966075 | 86920416 |
| (M.1 + M.0) + (H.0 + H.1) | 16 | 4127 | 2867892147 | 76102118 | 30 | 4025 | 2861902401 | 63087330 | 25 | 2789 | 2858870290 | 87638277 |
| (M.1 + M.0) + (H.1 + H.0) | 16 | 4116 | 2867893559 | 76102118 | 30 | 4027 | 2861965666 | 63087330 | 16 | 2785 | 2858947736 | 86920416 |

**Supplementary Table 6.** Selection of pseudo-molecules for each chromosome sequence for JG2

| **Chromosome** | **selected meta-scaffold** | **chromosome** | **selected meta-scaffold** |
| --- | --- | --- | --- |
| 1 | (jg1a + jg1c) + jg1b | 13 | jg1b + (jg1a + jg1c) |
| 2 | jg1a + (jg1c + jg1b) | 14 | jg1a + (jg1c + jg1b) |
| 3 | jg1c + (jg1a + jg1b) | 15 | jg1a +(jg1b + jg1c) |
| 4 | jg1a + (jg1c + jg1b) | 16 | (jg1c + jg1a) + jg1b |
| 5 | (jg1b + jg1a) + jg1c | 17 | jg1a + (jg1b + jg1c) |
| 6 | jg1c + (jg1a + jg1b) | 18 | jg1c + (jg1b + jg1a) |
| 7 | jg1c + (jg1b + jg1a) | 19 | jg1a + (jg1b + jg1c) |
| 8 | jg1b + (jg1c + jg1a) | 20 | (jg1a + jg1b) + jg1c |
| 9 | jg1b + (jg1c + jg1a) | 21 | (jg1b + jg1c) + jg1a |
| 10 | jg1b + (jg1a + jg1c) | 22 | (jg1c + jg1b) + jg1a |
| 11 | (jg1c + jg1b) + jg1a | X | (jg1a + jg1b) + jg1c |
| 12 | jg1c + (jg1a + jg1b) | Y | (jg1a + jg1b) + jg1c |
